# Untethered Muscle Tracking Using Magnetomicrometry

**DOI:** 10.1101/2022.08.02.502527

**Authors:** Cameron R. Taylor, Seong Ho Yeon, William H. Clark, Ellen G. Clarrissimeaux, Mary Kate O’Donnell, Thomas J. Roberts, Hugh M. Herr

## Abstract

Muscle tissue drives nearly all movement in the animal kingdom, providing power, mobility, and dexterity. Technologies for measuring muscle tissue motion, such as sonomicrometry, fluoromicrometry, and ultrasound, have significantly advanced our understanding of biomechanics. Yet, the field lacks the ability to monitor muscle tissue motion for animal behavior outside the lab. Towards addressing this issue, we previously introduced magnetomicrometry, a method that uses magnetic beads to wirelessly monitor muscle tissue length changes, and we validated magnetomicrometry via tightly-controlled in situ testing. In this study we validate the accuracy of magnetomicrometry against fluoromicrometry during untethered running in an in vivo turkey model. We demonstrate real-time muscle tissue length tracking of the freely-moving turkeys executing various motor activities, including ramp ascent and descent, vertical ascent and descent, and free roaming movement. Given the demonstrated capacity of magnetomicrometry to track muscle movement in untethered animals, we feel that this technique will enable new scientific explorations and an improved understanding of muscle function.

## Introduction

Muscle length measurements have driven important discoveries in movement biomechanics (Fowler et al. 1993), informed models of motor control (Roberts et al. 1997; Prilutsky et al. 1996), and provided strategies for prosthetic and robotic design (Eilenberg et al. 2010). For decades, sonomicrometry (SM) has informed how muscles move, providing high accuracy (70 μm resolution) and high bandwidth (>250 Hz) (Griffiths 1987). Fluoromicrometry (FM) expanded the muscle tracking toolkit, enabling high accuracy (90 μm precision) and high bandwidth (>250 Hz) for high-marker-count tracking (Brainerd et al. 2010; Camp et al. 2016). Further, image-based ultrasound (U/S) added the capability to non-invasively track muscle geometries (Fukunaga et al. 2001; Sikdar et al. 2014; Clark and Franz 2021).

Yet, collecting direct muscle length measurements in natural environments remains infeasible, and thus indirect muscle length estimation is still used for observing natural movements. For instance, muscle lengths are estimated using joint angles via biophysical models (Delp et al. 2007). These approximations are used due to the limitations of current muscle motion sensing techniques, all of which are tethered or bulky. SM and U/S both require tethered connections to bulky hardware for sensing (Biewener et al. 1998; Clark and Franz 2021), with SM requiring advanced surgery and percutaneous wires. And while FM does not require a tethered connection, it is limited to a volume approximately the size of a soccer ball, requires equipment the size of a small room, and is time-constrained due to thermal limitations and subject radiation exposure (Brainerd et al. 2010).

Present muscle length tracking technologies also require substantial post-processing time, hindering their use in longitudinal studies. SM requires accounting for and filtering out artifacts such as triggering errors (Marsh 2016), FM requires point labeling in stereo images (Brainerd et al. 2010), and U/S requires fascicle labeling (Van Hooren et al. 2020), all of which require at least some manual processing. While machine learning techniques have shown potential for automatic fascicle length tracking from ultrasound images, the current lack of reliability in tracking cross-activity measurements (*R*^2^=0.05 for single-subject cross-activity training of an SVM) prevents such a strategy from being applicable toward sensing fascicle lengths during natural movement (Rosa et al. 2021).

Researchers need a sensing platform that can operate untethered in natural environments, sensing the full dynamic range of muscle movement in context. To address this need, we developed magnetomicrometry (MM), a minimally-invasive strategy for portable, real-time muscle tracking. MM uses an array of magnetic field sensors to locate and calculate the distance between two implanted magnetic beads with sub-millisecond time delay. This distance provides a measurement of the muscle tissue length between the implanted beads. MM allows continuous recording over an indefinite collection interval extending across hours, with the potential for continuous use across days, weeks, or years.

In prior work, we validated the MM concept against FM via tightly controlled in situ tests (Taylor et al. 2021). However, it has not previously been empirically demonstrated that the MM technique is robust for recording during untethered locomotion. In the present study we address this question. We investigate the robustness of MM during untethered activity that exhibits soft tissue artifacts (i.e., movement of the magnetic field sensors relative to the muscle) and changes in the relative orientation of the ambient magnetic field.

Herein we present MM as a robust, practical, and effective strategy for measuring muscle tissue length in an untethered freely-moving animal model. We first apply this technique to turkeys running on a treadmill and compare MM to FM to determine the method’s accuracy. We then further investigate the use of MM to track muscle tissue length in freely-moving animals during ramp ascent and descent, vertical ascent and descent, and free roaming movement. We hypothesize that muscle tissue lengths during untethered motion can be tracked via MM with submillimeter accuracy and a strong correlation (R^2^ > 0.5) to FM. Our validation of this tool in a mobile context enables tracking and investigation of muscle physiology in settings previously inaccessible to biomechanics researchers.

## Results

### Accuracy Validation of Magnetomicrometry Against Fluoromicrometry

To verify MM tracking accuracy during untethered activity, we tracked implanted magnetic bead pairs in turkey gastrocnemius muscles (right leg, three turkeys) using both MM and FM while the turkeys walked and ran at multiple speeds on a treadmill (see Figure 1 for the setup and tracking results, see Supplementary Figure 1 for a 3-D scan of the MM sensing array).

**Figure 1:**
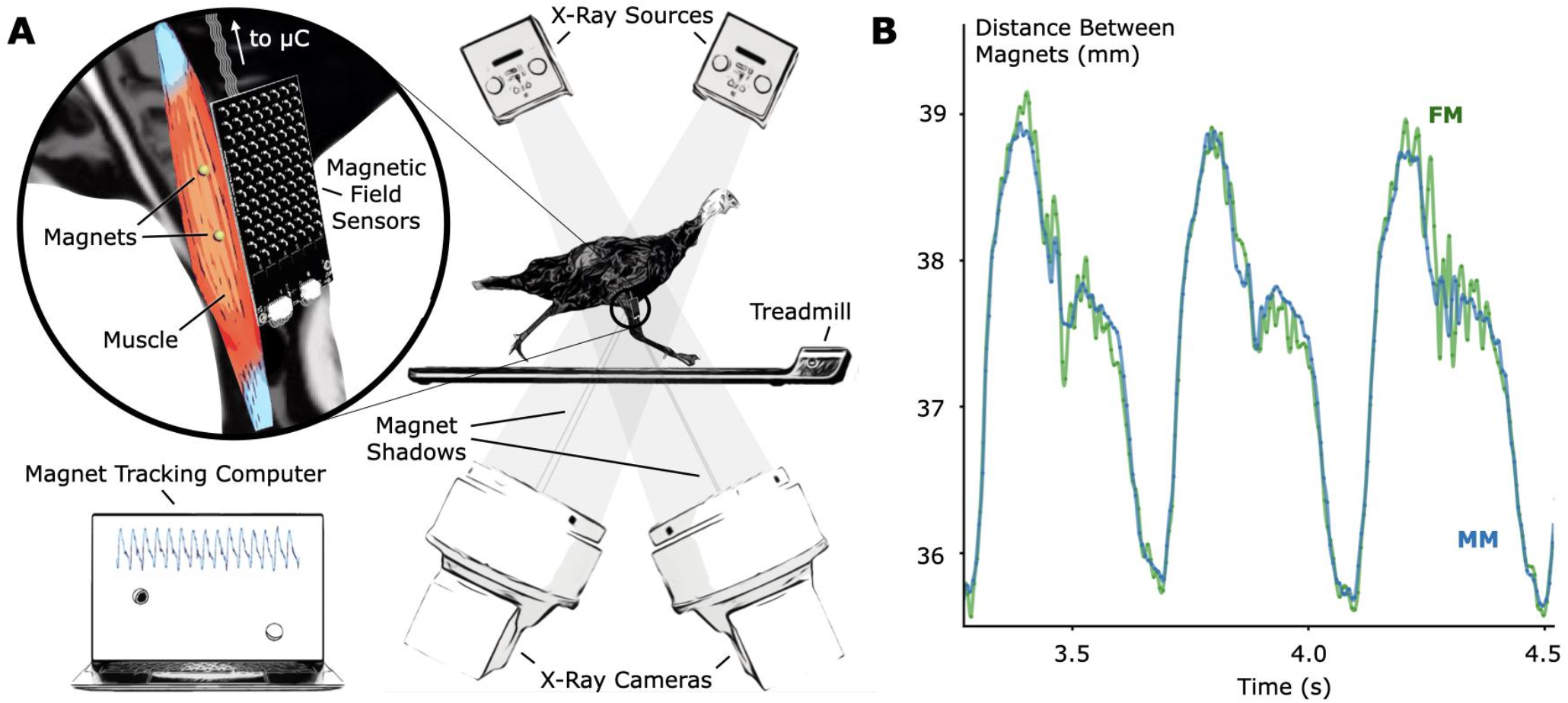
Validation of Untethered Muscle Tracking using Magnetomicrometry. **(A)** A magnetic field sensing array on the surface of the leg tracks the positions of two magnetic beads implanted into the muscle. A feather microcontroller (μC) in the turkey feathers wirelessly transmits the magnetic field data to a magnet tracking computer that calculates and displays the magnetomicrometry (MM) signal in real time. The turkeys walked and ran on a treadmill while x-ray video cameras recorded synchronized fluoromicrometry (FM) data for post-processing. **(B)** Comparison of MM (blue) with FM (green) to validate the MM accuracy. These representative results during running gait show the submillimeter accuracy of MM during untethered muscle length tracking.

We compared the distances between the magnetic bead positions as measured by MM with their distances as measured by FM to evaluate accuracy during the treadmill activity (see Figure 2). The coefficients of determination (*R^2^* values) between MM and FM were 0.952, 0.860, and 0.967 for Birds A, B, and C, respectively (see also Supplementary Figure 2). The differences between MM and FM were −0.099 ± 0.186 mm, −0.526 ± 0.298 mm, and −0.546 ± 0.184 mm for Birds A, B, and C, respectively (see Supplementary Figure 3).

**Figure 2:**
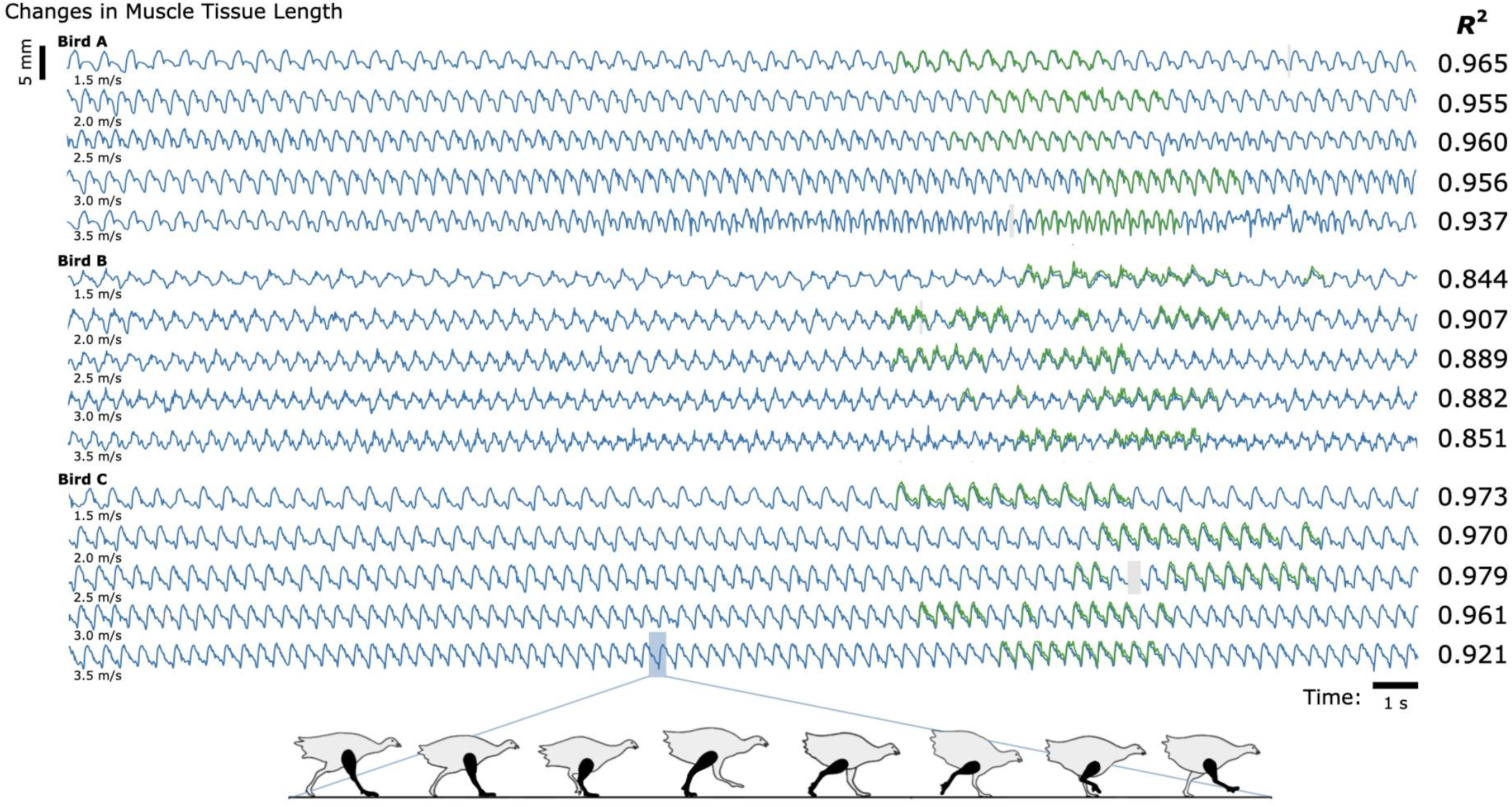
Untethered Muscle Tracking During Treadmill Running: Magnetomicrometry Versus Fluoromicrometry. Changes in muscle tissue length measured by MM (blue) and FM (green) for three turkeys at five speeds (30 s shown for each speed). The column to the right of the plots gives the coefficients of determination (R^2^) between magnetomicrometry and fluoromicrometry corresponding to each turkey and speed. Gaps in the fluoromicrometry data are due to researcher selection of full gait cycles during which both magnetic beads were visible in both x-ray images. Gaps in the magnetomicrometry data (gray) are due to packet drops during wireless transmission of the magnetic field signals to the tracking computer (gaps below 50 ms interpolated in gray, gaps above 50 ms highlighted in gray). The turkey gait diagram below the plots shows the corresponding gait phases over one gait cycle.

To determine the study-specific reliability of the manual FM processing (marker position labeling in the X-ray video data), ten gait cycles of raw FM data were independently manually relabeled three times for one bird at one speed. Across these three labelings for these ten gait cycles, manual FM processing was consistent to a standard deviation of 0.098 mm (see Supplementary Figure 4 for more details).

MM’s 99-th percentile tracking time delays were 0.698 ms, 0.690 ms, and 0.664 ms for Birds A, B, and C, respectively (see also Supplementary Figure 5), and the MM data did not require any post-processing. In contrast, post-processing the FM data into marker-to-marker distances required approximately 84 manual processing hours spread across multiple months.

### Untethered Muscle Tracking Across Various Activities

To investigate the feasibility of using MM during dynamic, natural motion, we constructed a series of obstacles for the turkeys to navigate. Specifically, we provided the turkeys with two ramp inclines (10° and 18°, see Figure 3) and three vertical elevation changes (20 cm, 41 cm, and 61 cm, see Figure 4). Because the purpose of these activities was to explore the range of dynamic motions that could be captured, we did not train the birds to navigate the ramps or vertical elevation changes repetitively, and thus variability is expected within the repeated tasks.

**Figure 3:**
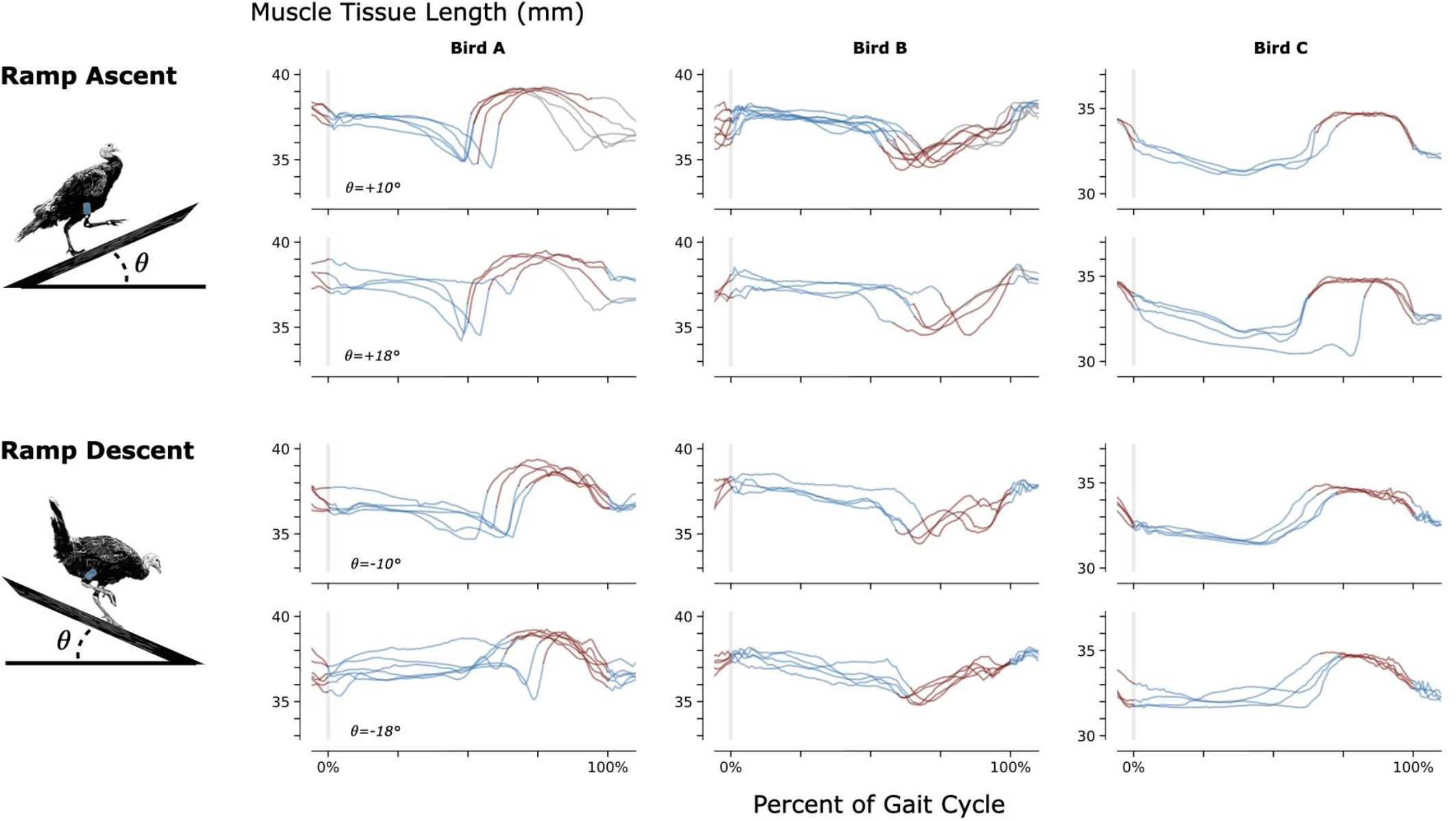
Muscle Tissue Length During Non-Synchronous Ramp Ascent and Descent. We used magnetomicrometry to track muscle tissue length during ramp ascent and descent at two inclines for all three birds. Data for each bird and each slope are synchronized at right leg toe strike (indicated by the vertical gray line) and normalized from toe strike to toe strike. Variability between curves reflects gait cycle variability during untrained ramp navigation. Muscle tissue length is plotted in blue for right leg stance, in red for right leg swing, and in gray where video did not allow gait-phase labeling. We recorded at least three gait cycles of each activity for each bird.

**Figure 4:**
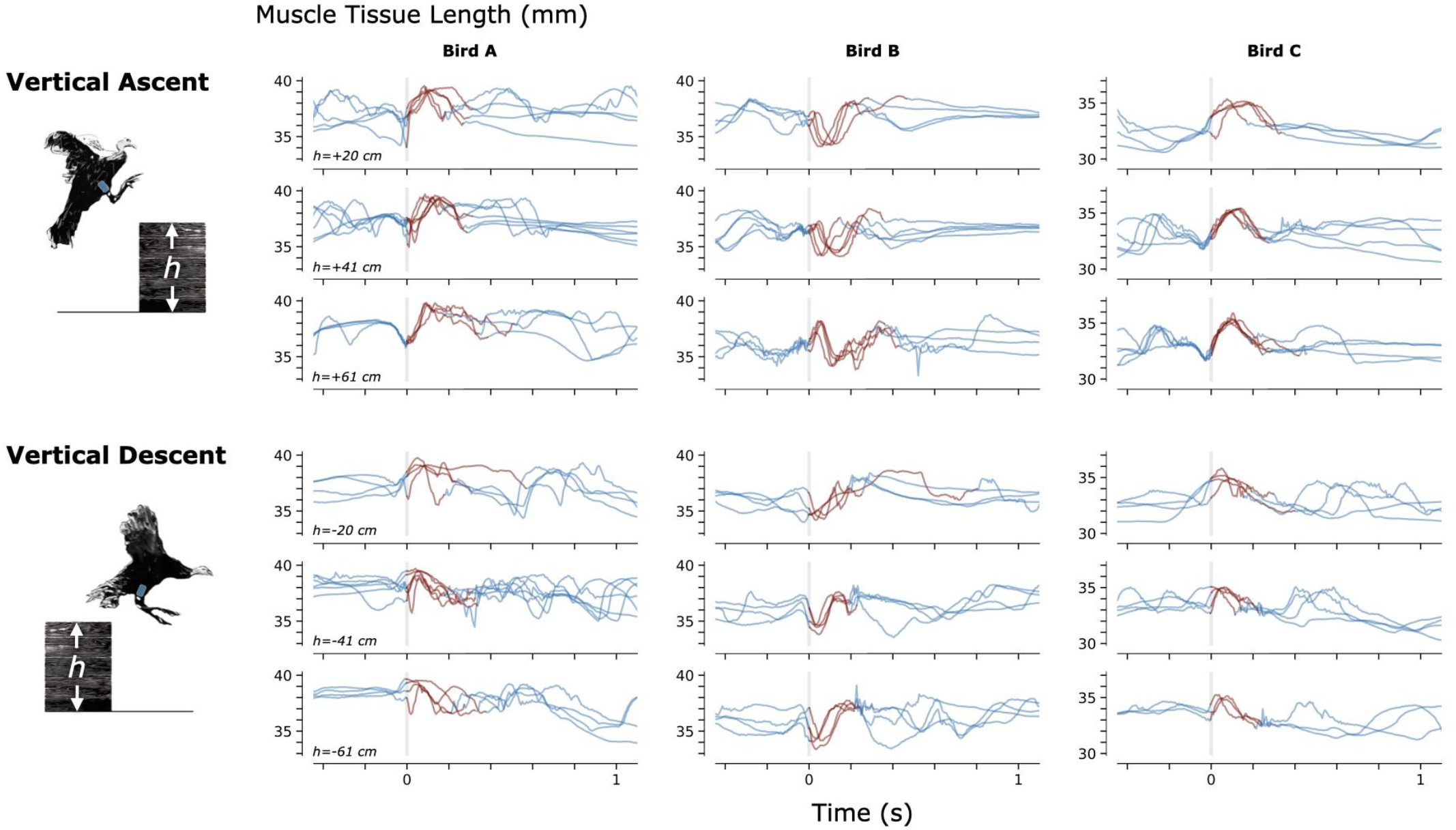
Muscle Tissue Length During Non-Synchronous Vertical Ascent and Descent. We used magnetomicrometry to track muscle tissue length during vertical ascent and descent at three heights for all three birds. Data for each bird and each height are synchronized at right leg toe-off (start of the aerial phase, indicated by the vertical gray line). Variability between curves reflects movement variability during untrained vertical ascent and descent. Muscle tissue length during contact with the ground is plotted in blue, and muscle tissue length during the aerial phase is plotted in red. All data are shown, including scenarios in which significant wing-flapping occurred during jump up or down. We captured at least three recordings of each activity for each bird.

To further validate the accuracy of MM used during navigation of ramps and vertical elevation changes, we analyzed the magnetic bead tracking data from these activities to find the range of the tracked three-dimensional magnetic bead positions (see Supplementary Figure 6). We then affixed two magnetic beads 40 mm apart, validated the distance between them using FM (40.000 ± 0.017 mm), and swept this FM-validated magnetic bead pair under the MM sensing array through a volume exceeding these ranges (see Supplementary Figure 7). We monitored deviations from 40 mm in the MM signal during these benchtop tests and found a 99-th percentile error (e99%) of 1.000 mm (rounded up to the nearest micrometer).

Finally, to explore whether untethered muscle tracking via MM is viable in a fully free roaming context, we tracked muscle tissue length while one turkey (Bird A) roamed freely about its enclosure. The results of this data collection are shown in Figure 5.

**Figure 5:**
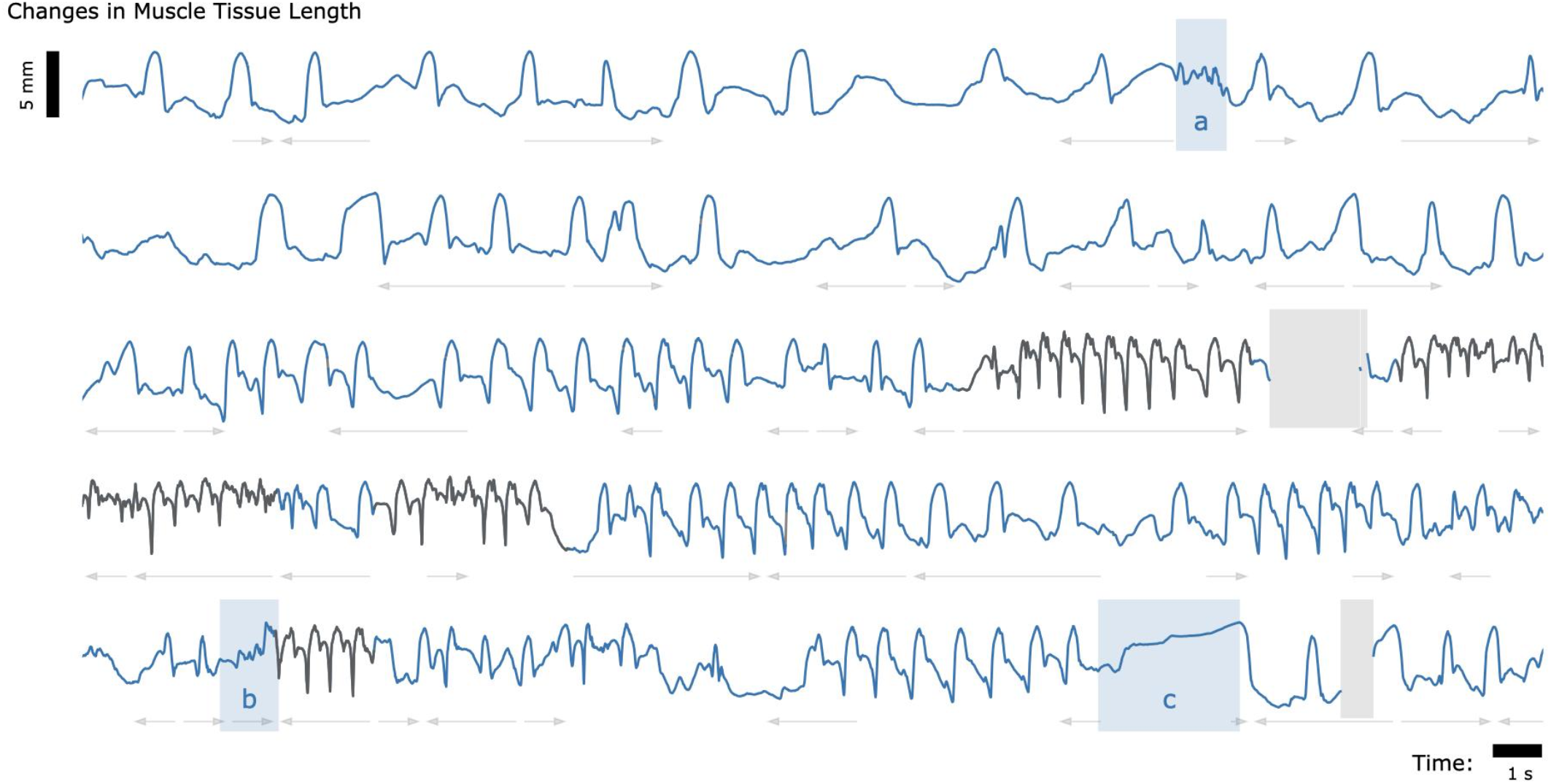
Muscle Tissue Length During Free Roaming Movement. Magnetomicrometry data was continuously collected for 150 seconds during free roaming activity. Muscle tissue length is plotted in blue during standing and walking and plotted in black during running. Blue highlighted regions indicate muscle tissue length during (a) feather ruffling, (b) jumping, and (c) balancing on one leg. Gray arrows indicate when the turkey was turning left (left arrows) or turning right (right arrows). Gaps due to wireless transmission packet drops are shown in gray, as described in Figure 2.

## Discussion

We find that MM enables untethered muscle tissue length tracking with high correlation to FM (R^2^ of 0.952, 0.860, and 0.967 for Birds A, B, and C, respectively) and submillimeter accuracy (AVG±SD of −0.099 ± 0.186 mm, −0.526 ± 0.298 mm, and −0.546 ± 0.184 mm for Birds A, B, and C, respectively). These findings enable tracking and investigation of muscle contractile behavior in settings previously inaccessible to biomechanics researchers.

### Accuracy Validation

The standard we used here to assess the accuracy of muscle length tracking using MM was FM. For magnets implanted superficially in muscles (at depths less than 2 cm), MM exhibits less noise than FM, but FM has the advantage of higher accuracy, especially at greater tissue depths (tracking depths in this study ranged from 11.2 mm to 26.6 mm). Indeed, our tests showed that for unobscured markers moving through the X-ray volume, FM was accurate to 0.030 mm. However, we note that marker tracking noise was a challenge for FM in this particular study due to the use of a large animal and the presence of hardware (the MM sensing array) that regularly obscured the markers during the tracking. These factors resulted in substantial manual labeling noise in the FM signal of 0.098 mm, instead of the 0.030 mm noise we found in our FM accuracy test, affecting the accuracy standard deviations reported above. Accounting for this manual labeling noise gives adjusted accuracy standard deviations of 0.158, 0.281, and 0.156 mm, for Birds A, B, and C, respectively (see Supplementary Figure 3).

Constraints to imaging volume make FM impractical during large-animal variable terrain activity, so we performed retrospective benchtop accuracy testing to further validate the MM data collected during navigation of the ramps and vertical elevation changes (see Supplementary Figure 7). The error we observed in the benchtop tests (e_99%_ < 1 mm) was acceptable in comparison with the magnitude of the muscle contractions we observed during the variable terrain activity (average MM signal magnitude was 4.5 mm peak-to-peak). This suggests that MM robustly tracked the muscle tissue lengths during the variable terrain activities, despite any soft tissue artifacts that may have occurred during the dynamic movements required by those activities. These tests, however, highlight the importance of sensor placement. Higher accuracy is achieved when the MM sensing array is properly placed – centered over the implanted beads. MM with perfect magnetic field sensing would, in theory, be unaffected by movement of the board relative to the implanted beads, but the errors we observed suggest that the sensors are nonlinear. Magnet tracking nonlinearity compensation (e.g., via sensor calibration or three-dimensional sensor geometries) is thus an important area for future research. Meanwhile, in future work, larger sensing arrays with broader coverage would be advantageous to mitigate the need for careful placement of the array.

### Ambient Magnetic Fields

The software-based magnetic disturbance compensation we employed here (Taylor et al. 2019) was sufficient to compensate for ambient magnetic fields during untethered muscle tracking in the presence of large hydraulic ferromagnetic lift tables, a large, active treadmill motor, and a room full of active X-ray equipment. However, our uniform disturbance compensation strategy may be insufficient for the exceptional situation where a large ferromagnetic object is immediately adjacent to (within a few centimeters of) the tracked muscle. Thus, software-based compensation for spatially-non-uniform ambient magnetic fields may still be a valuable direction for future work to extend the robustness of MM to that potential scenario. Alternatively, ferromagnetic shielding could be used to physically perform disturbance compensation (Tarantino et al. 2017), but the shield would need to be sufficiently far away to prevent it from acting like a magnetic mirror, creating “image” magnets that would need to be tracked as well (Hammond 1960). Further, effective shielding would need to be thick enough to redirect most or all magnetic field disturbances, presenting a trade-off between the weight and the efficacy of the shielding.

### Range of Behaviors

Figures 3, 4, and 5 provide a sample of the range of behaviors that can be tracked using magnetomicrometry. Consistency in the curves was not strived for, expected, or desired. Rather, we intentionally preserved anomalous events in those data, such as single or multiple wing flaps during vertical ascent and descent and variable speed during ramp navigation, to explore the range of motor activities during which we could track the muscle activity.

### Applications

MM has the potential to work across scales (see Figure 6), from the ability to track both full-body and muscle movement of small organisms to the ability to track large magnetic beads implanted deep into large animal models. Mathematically, if the number of sensors is fixed and all system dimensions are scaled, the error as a percent of scaled magnetic bead excursion will remain unchanged (Taylor et al. 2019). However, larger sensing arrays can in principle be used when tracking very small or very large animals, resulting in an increase in tracking accuracy at those extremes. For instance, when tracking small animal muscle tissue, additional sensors can be embedded into the animal’s environment, and when tracking large animal muscle tissue, the increased animal size accommodates the mounting of additional sensors to the animal. Thus, context-specific magnetic bead tracking systems can take advantage of the unique geometries afforded at each scale.

**Figure 6:**
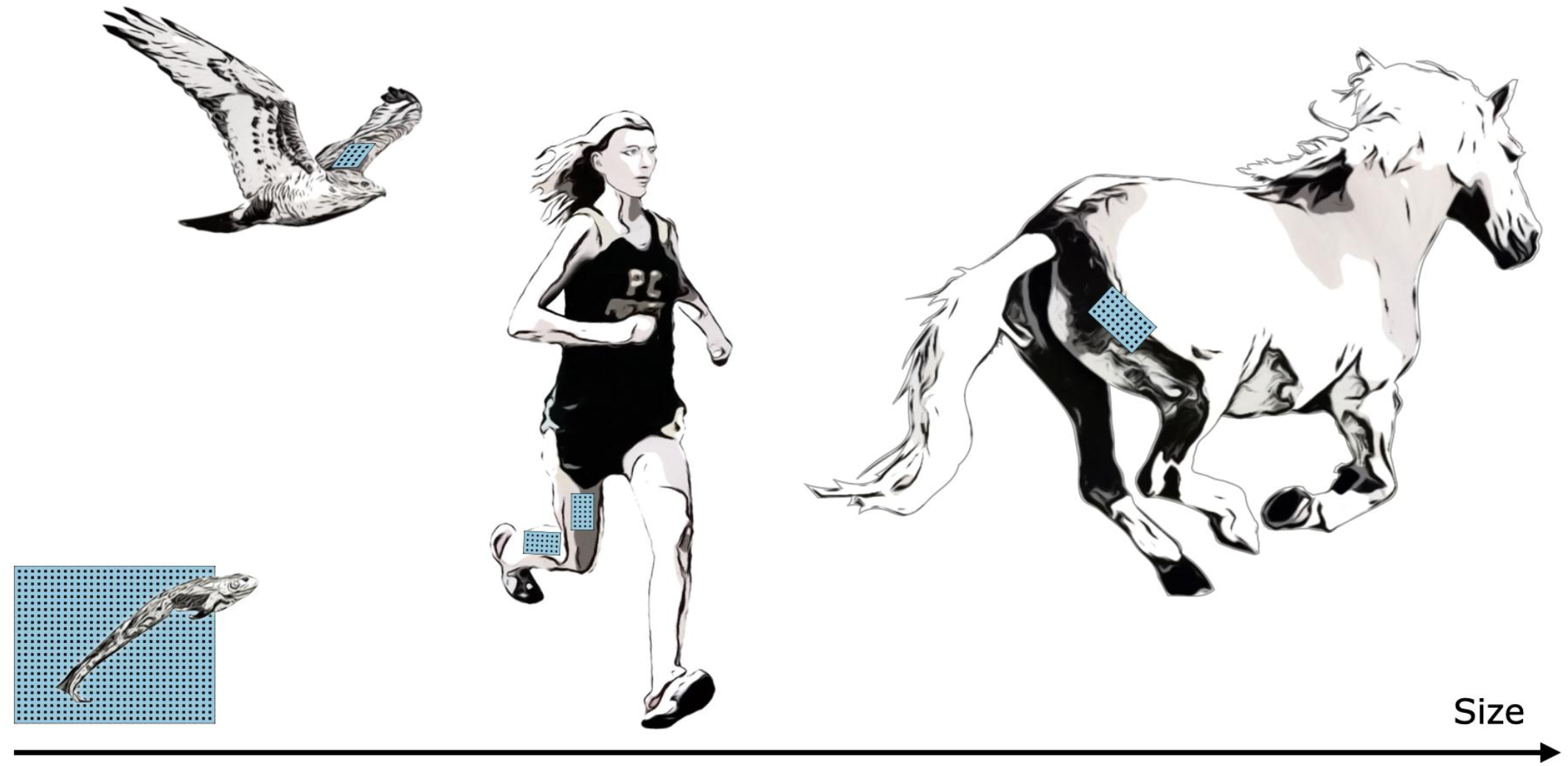
Muscle Tracking Across Scales. By changing the size of the magnetic field sensing array, we can track the distance between magnets at closer or farther distances, allowing us in principle to track muscle tissues at a range of scales, including frogs, hawks, persons, horses, or other animals. For small animals, such as the frog shown at bottom left, a fixed array below or beside the animal could track both the position of the animal and the muscle tissue length.

Not only can MM be used across size scales, but across time scales as well. Because there is no need for post-processing, MM data can be collected continuously, enabling the potential for longitudinal studies, including investigations into mechanisms such as neural degradation or plasticity over time.

MM’s muscle length and velocity signals are different from, and complementary to, the signals from electromyography (EMG). While EMG provides a measure of muscle activation, which results more directly from neural commands, muscle length and velocity give information on the shape of the muscle, which in turn can refine our understanding of muscle physiology during a given motion task. Indeed, the combination of MM and EMG will allow for increased physiological understanding in new contexts where animals are in their natural environments.

In parallel work, we also demonstrate the viability of magnetic bead implants for human use, verifying comfort, lack of implant migration, and biocompatibility (Taylor et al. 2022). Due to the untethered nature of MM, this technique has applications in prosthetic and exoskeletal control. In its most straightforward implementation, a motor controller could directly control a robotic joint using the distance between two beads in each muscle of a flexor-extensor pair. However, the ability for MM to track additional muscles and to work in combination with EMG enables a range of new strategies for human-machine interfacing.

### Limitations

In this study, we implanted the magnetic beads approximately 3.5 cm away from one another, based on previous work (Taylor et al. 2021), to ensure that the magnetic beads would not migrate toward one another. If smaller or larger (or differently shaped) magnetic bead implants are used (for instance, in a smaller or larger animal model), the effect of separation distance on stability against migration would need to be re-investigated for the different sizes (and different magnetization strengths) of the implants.

For the benchtop accuracy validation tests, we assumed that the tracked bead positions were a good approximation for the true bead positions. We used the magnetic bead position tracking information from the variable terrain MM data to determine the boundaries of the volume to test. Then, at the start of the tests, we used tracked bead positions to locate the centered, minimum-depth position (the closest position within the full scale range of the sensors), then used blocks of known dimensions to sweep through the benchtop-emulated tissue depth and enforce the volume boundaries. Noting that MM was accurate to within a millimeter throughout the volume, we found these assumptions reasonable for these tests.

As for any muscle tissue tracking, the location of the implanted tracking devices will determine the length measured. For studies where the aim is to relate measured length changes to muscle contractile properties (e.g., length-tension or force-velocity relationships), it is essential that the markers are aligned along the fascicle axis. In the present study we embedded magnets at locations in the turkey muscles that would ensure the magnets stayed in place over a period of months, and at depths that were favorable for sensor function. Thus, patterns of length change do not directly represent patterns of muscle fascicle length change and can be influenced significantly by dynamic changes in muscle architecture during contraction. This is reflected in the opposite muscle tissue length changes seen during the swing phase of Bird B relative to Birds A and C during ramp navigation (see Figure 3). MM, FM, and SM all suffer from this same issue, and thus for any of these techniques, careful surgical placement is warranted.

### Sensing Improvements

The suite of electronics for MM is immediately upgradeable as new industry standards develop. The tracking system benefits from global developments in low-cost magnetic field sensors due to the widespread manufacturing of inertial measurement units for devices such as cell phones, video game controllers, and autonomous vehicles. Continuing improvements in magnetic field sensors, capacitors, and microcontrollers will cause direct improvements to the accuracy, efficiency and speed of the tracking system and will allow the tracking of even smaller implants at greater depths.

### Summary

Here, we demonstrate the use of MM for untethered muscle tracking. We validate, against FM, the submillimeter accuracy of MM in an awake, active turkey model (R^2^ ≥ 0.860, *μ* ≤ 0.546 mm, *σ* ≤ 0.298 mm) with a real-time computing time delay of less than a millisecond (η_0.99_ ≤ 0.698 ms). We further demonstrate the use of MM in untethered muscle tracking during ramp ascent and descent, vertical ascent and descent, and free roaming movement. These results encourage the use of MM in future biomechanics investigations, as well as in prosthetic and exoskeletal control. We hope that MM will enable a variety of new experiments and technologies, and we look forward to the further development and application of this technology.

## Methods

All animal experiments were approved by the Institutional Animal Care and Use Committees at Brown University and the Massachusetts Institute of Technology. Wild turkeys (*Meleagris gallopavo*, adult female) were obtained from local breeders and maintained in the Animal Care Facility at Brown University on an ad libitum water and poultry feed diet. We used three animals in this study.

### Surgical Procedure

One pair of 3-mm-diameter Parylene-coated magnetic beads (N48SH) were implanted into the right lateral gastrocnemius muscle of each turkey, with a target magnetic bead separation distance of 3.5 cm. For details on the surgical procedure and implants, see Taylor et al. 2022. A one-month recovery period was given before the start of the data collection.

### Magnetomicrometry

For this study, we designed a custom magnetic field sensing array (see Figure 7). The sensing array was equipped with 96 magnetic field sensors (LIS3MDL, STMicroelectronics) spaced 5.08 mm apart in an 8-by-12 grid. Each sensor was supplied with nonmagnetic capacitors (VJ1206Y105KCXAT and VJ0603Y104KCXAT, Vishay). Seven digital multiplexers on the sensing array allowed time-domain multiplexing (one 74HC138BQ,115 multiplexing into six 74HC154BQ,118, Nexperia) via a wired connection. The sensing array was connected through a custom adapter board to an off-the-shelf wireless microcontroller embedded system (Feather M0 WiFi microcontroller, Adafruit), which was powered by a lithium-ion polymer battery (3.7 V, 1800 mA·h, 29 g). The microcontroller sampled the magnetic field signals at 155 Hz and wirelessly transmitted them to the magnet tracking computer via a WiFi router (Nighthawk R6900P, Netgear). The tracking algorithm ran in real-time on the magnet tracking computer, a Dell Precision 5550 laptop (Ubuntu 20.04 operating system) with 64 GB of random-access memory and an Intel i7 8-Core Processor, running at 2.30 GHz. The tracking algorithm used, including the strategy for disturbance compensation, is fully-detailed in previous work (Taylor et al. 2019).

**Figure 7:**
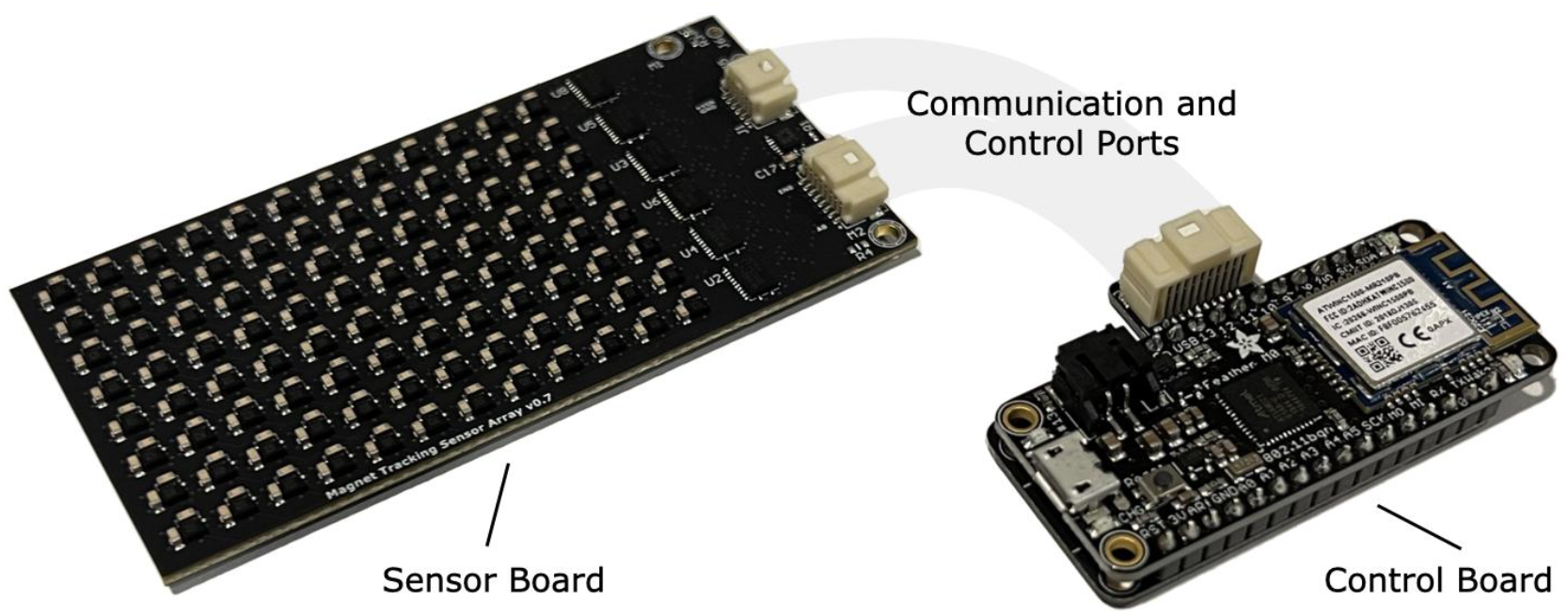
Magnetomicrometry Embedded System. We fabricated a custom sensor board (left) and a custom control board (right) for this study. The sensor board holds the magnetomicrometry sensing array, consisting of 96 magnetic field sensors arranged with a spacing of 5.08 mm. Digital multiplexers on the sensor board allow time-domain multiplexing, enabling a single microcontroller on the control board to communicate with and control all magnetic field sensors on the sensor board. The control board merges the data from the sensor board and streams the data wirelessly to the magnet tracking computer. The sensor board and control board weigh 24 g and 12 g, respectively.

We affixed the sensing array to the limb using Opsite Flexifix adhesive film (Smith & Nephew). To secure the array, we first applied a base layer of the adhesive film to the skin. We then positioned the sensing array over the leg and wrapped the adhesive film around the sensing array and the leg. To maintain a sufficient minimum distance between the magnetic beads and the sensing array, we positioned layers of foam between the base adhesive and the array. We then secured the control board and battery within the back feathers of the turkey.

### Accuracy Validation of Magnetomicrometry Against Fluoromicrometry

We used the W. M. Keck Foundation XROMM Facility at Brown University (Brainerd et al. 2010) to perform FM. We collected X-ray video from two intersecting X-ray beams oriented at 51 degrees relative to one another. We mounted a treadmill with a wooden base (TM145, Horizon Fitness) between the X-ray cameras and the X-ray sources and built a housing over the treadmill with a movable wall to position the birds within the capture window.

The turkeys walked and ran at five speeds (1.5, 2.0, 2.5, 3.0, and 3.5 m/s) in a randomized order until at least ten gait cycles were visible within the FM capture volume for each speed. We collected FM data at 155 Hz.

Time syncing was performed via a coaxial cable connection from FM to an off-the-shelf microcontroller development board (Teensy 4.1, Adafruit). The time-syncing microcontroller relayed the time sync signal to the magnet tracking computer via a custom adapter board.

### Untethered Muscle Tracking Across Various Activities

We constructed a hallway to guide the turkeys through the variable terrain activities. We stacked plyometric boxes (Yes4All) in the hallway to heights of 20 cm, 41 cm, and 61 cm for vertical ascent and descent. Upon completion of the vertical ascent and descent tests, we placed ramps (Happy Ride Folding Dog Ramp, PetSafe) up to and down from the plyometric boxes at inclines of 10° and 18° for the turkeys to ascend and descend. Separate from the variable terrain activities, we then allowed one turkey (Bird A) to roam freely within its enclosure while we continued to record MM.

### Benchtop Magnetomicrometry Validation for the Variable Terrain Activity

For benchtop testing, we used super glue (Krazy Glue) to affix each of two N48SH magnetic beads into two 1×1 round LEGO plates. We attached these round LEGO plates to a 1×6 LEGO technic block, one at each end, to separate the pair of magnetic beads by a fixed distance of 40 mm (see Supplementary Figure 7A). We imaged this pair of beads using FM to validate the 40 mm fixed distance between them. Specifically, after the MM accuracy validation of two of the birds, we collected FM data with the magnet pair statically in the volume and used the average of the last three seconds from each of these two FM collections to confirm the distance between the two beads.

To determine the volume to sweep this FM-validated 40-mm-distanced bead pair during benchtop testing, we analyzed the magnetic bead tracking data from all ramps and vertical elevation changes to determine the full range of the three-dimensional magnetic bead positions relative to the MM sensing array across all three turkeys.

We first aligned and centered the magnet pair under the array, and we used the tracked magnet z-positions to place the magnets at the closest depth that was possible while still within the full-scale sensing range of the sensors (~1 cm). We then used 3.2-mm-thick 1×6 LEGO plates to enforce the remaining depths (see Supplementary Figure 7A). At each depth, we manually swept from center out and back along the x and y axes to the point where the farthest magnet reached just beyond the test volume requirements derived from the variable terrain activity (see Supplementary Figure 7B).

### Data Analysis

We post-processed the FM data using XMA Lab (Knörlein et al. 2016). All FM and MM data were left unfiltered.

We aligned the MM and FM data using the time sync signal and linearly interpolated the FM data at the MM measurement time points. Then, due to imprecision of the time sync signal from the X-ray system, we used local optimization to further align the MM and FM signals while iteratively interpolating FM. During one trial (one data collection for one turkey at one speed), where the tracking computer did not receive a time sync, we used global optimization to align the MM and FM signals. To validate the use of global optimization for synchronization, we tested this same global optimization on all other trials and found that the global optimization successfully located all time sync signals.

We estimated the noise from manual FM processing by independently processing one set of ten gait cycles three times (manually re-processing the video data twice without reference to the previously processed data). We then calculated the variance at each time point and used the square root of the average variance as our estimate of the FM manual processing noise (see Supplementary Figure 4). We calculated the adjusted MM noise by subtracting the average variance of the FM manual processing noise from the variance of the difference between the MM and FM signals for each bird, then taking the square root.

In Figure 3, for gait cycles where only one toe strike was visible in the video, we normalized the gait cycle using the timing of the peak MM signals in the previous and current gait cycles.

In Supplementary Figure 7, all data are shown plotted as a scatterplot with a smoothing cubic spline approximation, using a smoothing parameter of 0.8 (de Boor 1978). The plotted standard deviation was calculated as the root-mean-square of the spline-adjusted values, also smoothed with a smoothing parameter of 0.8.

## Author contributions

CT developed the magnetomicrometry strategy, led the experimental conception and design, assisted in surgeries, performed data collection, documentation, and analysis, and led the manuscript preparation. SY designed, validated, and oversaw the fabrication of the magnetic field sensing embedded system, set up the WiFi-enabled real-time measurement framework, contributed to experimental conception and design, assisted in surgeries, performed data collection, documentation, and analysis, and contributed to manuscript preparation. WC, EC contributed to experimental conception and design, assisted in surgeries, performed data collection, documentation, and analysis, and contributed to manuscript preparation. MO contributed to experimental conception and design, assisted in surgeries, and contributed to manuscript preparation. TR contributed to experimental conception and design, led the surgeries, assisted in data collection and analysis, and contributed to manuscript preparation. HH conceived the magnetomicrometry strategy, oversaw project funding, assisted with general study management, contributed to experimental conception and design, assisted in data analysis, and aided in manuscript preparation.

## Acknowledgments

The authors thank, non-exhaustively, Elizabeth Brainerd, Andreas Burger, Ziel Camara, John Capano, Matthew Carty, Tyler Clites, Charlene Condon, Kathy Cormier, Jimmy Day, Bruce Deffenbaugh, Jennie Ehlert, Wendy Ehlert, Rachel Fleming, Lisa Freed, Robert Gnos, Deborah Grayeski, Alan Grodzinsky, Ayse Guvenilir, Kale Hansen, Mallory Hansen, Guillermo Herrera-Arcos, Crystal Jones, Kylie Kelley, Duncan Lee, Aimee Liu, Richard Marsh, Richard Molin, Chad Munro, Jarrod Petersen, Mitchel Resnick, Lindsey Reynolds, Andy Robinson, Jacob Rose, Amy Rutter, Shriya Srinivasan, Erika Tavares, Sara Taylor and Beni Winet for their helpful advice, suggestions, feedback, and support. Inclusion in this list of acknowledgments does not indicate endorsement of this work.

## Data Availability

All data needed to support the conclusions in the paper are included in the main text and Supplementary Materials. In addition, the raw magnetomicrometry and processed fluoromicrometry data is available via the following Dropbox link: https://www.dropbox.com/s/qws2wkv9bh9uyrv/Untethered_Muscle_Tracking_Supplementary_Materials.zip?dl=0.

---

**Supplementary Figure 1:**
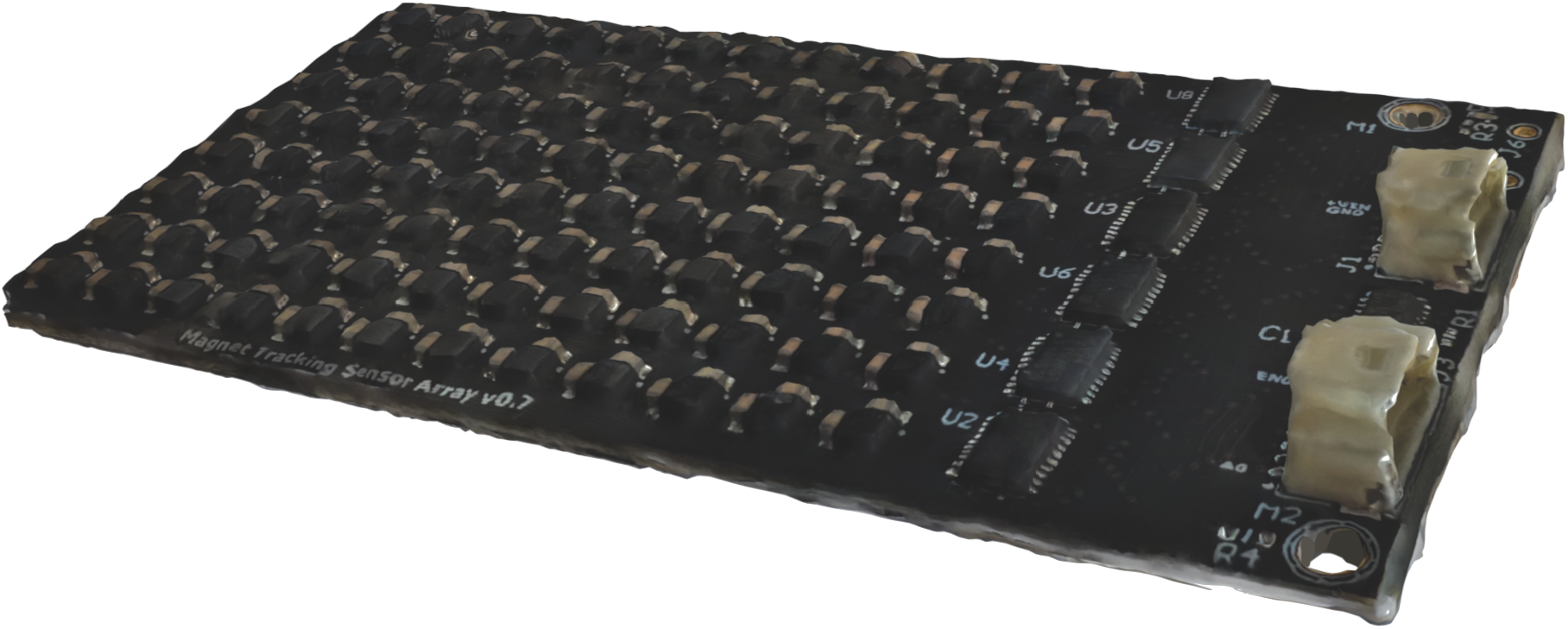
3D Scan of the Magnetomicrometry Sensor Board. See supplementary folder 3_D_Scan_of_MM_Sensor_Board for a 3-D scan of the magnetomicrometry sensor board in .obj format with associated reference files (.mtl, .png).

**Supplementary Figure 2:**
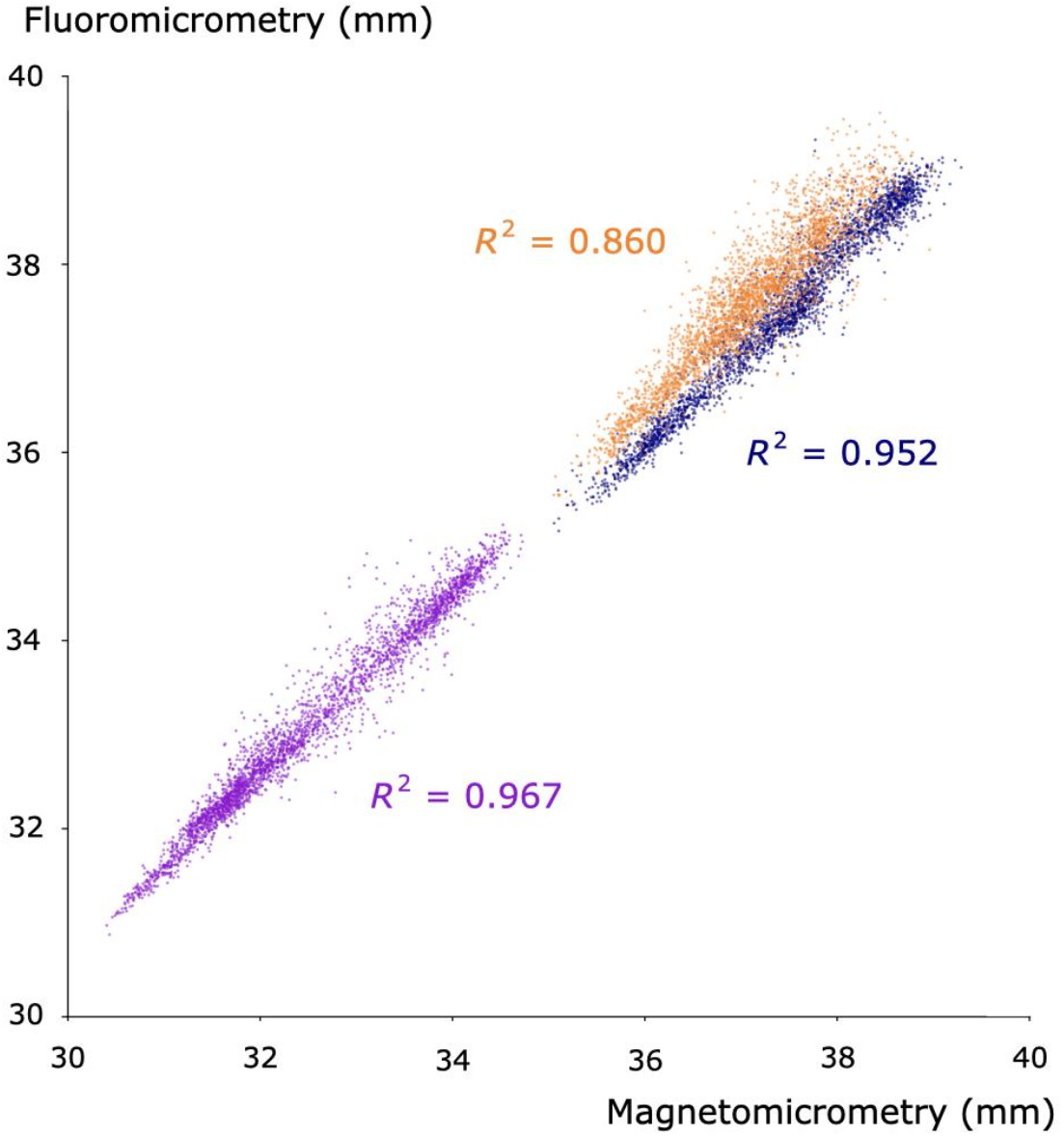
Coefficients of Determination (R^2^ values) between Magnetomicrometry and Fluoromicrometry. We compared all magnetomicrometry measurements (horizontal axis) across 50 gait cycles of turkey running (10 gait cycles at each of 5 speeds, for each bird) against time-synchronized, interpolated fluoromicrometry measurements (vertical axis). Data are plotted in blue, orange, and purple for Birds A, B, and C, respectively. Coefficients of determination (R^2^ values, shown in corresponding colors) for each bird were 0.952, 0.860, and 0.967, respectively.

**Supplementary Figure 3:**
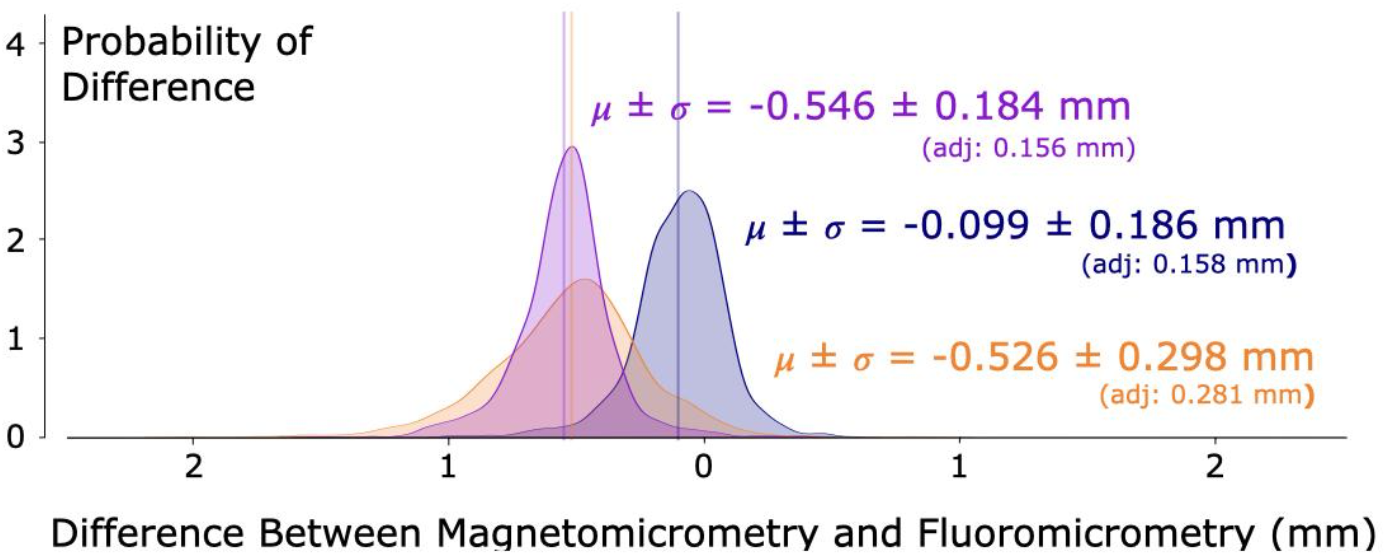
Kernel Density Estimates of Differences between Magnetomicrometry and Fluoromicrometry. We subtracted time-synchronized, interpolated fluoromicrometry measurements from magnetomicrometry measurements across all 50 gait cycles of turkey running (10 gait cycles at each of 5 speeds, for each bird). Kernel density estimates show the distribution (vertical axis) of these differences (horizontal axis), with data plotted in blue, orange, and purple for Birds A, B, and C, respectively. Vertical lines indicate mean offsets (in corresponding colors), with the mean offsets and standard deviations labeled. Adjusted standard deviations that compensate for fluoromicrometry noise (0.098 mm, standard deviation) indicate an estimate of the magnetomicrometry measurement noise.

**Supplementary Figure 4:**
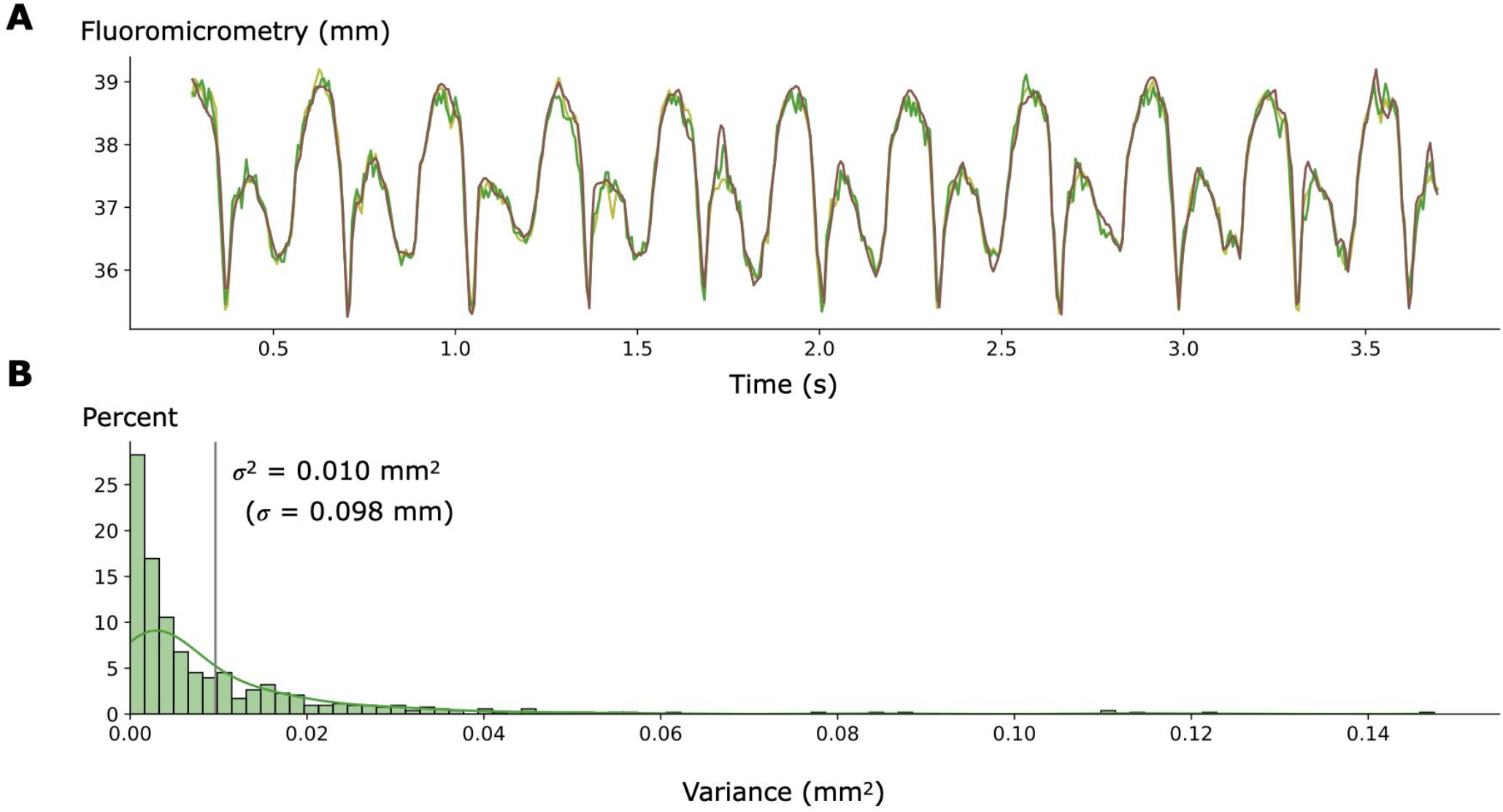
Study-Specific Limitations to the Use of Fluoromicrometry. **(A)** Ten gait cycles of raw fluoromicrometry video data (Bird A, 3.5 m/s), independently manually re-labeled three times, are shown in light green, dark green, and brown. **(B)** The histogram shows the distribution (vertical axis) of the variance (horizontal axis) between the three manual processed fluoromicrometry signals at each timepoint of the data, with a kernel density estimation curve overlay. A vertical line indicates the average variance, 0.010 mm^2, and the corresponding square root of the variance is 0.098 mm.

**Supplementary Figure 5:**
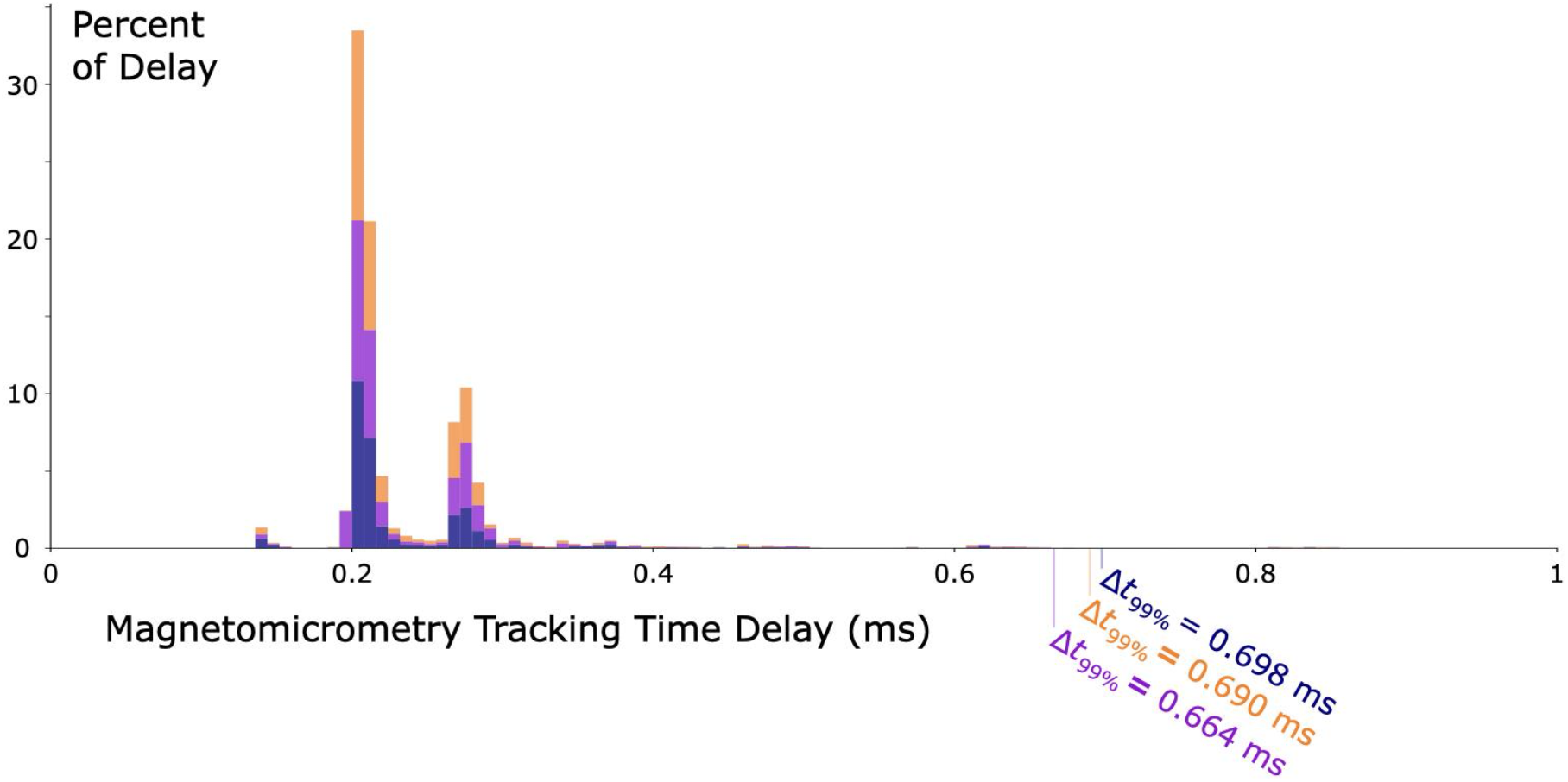
Magnetomicrometry Tracking Time Delay. The magnet tracking computer recorded the times it received the magnetic field data and the times it completed the magnet tracking algorithm. The distribution (vertical axis) of the difference between these times (horizontal axis) is the tracking time delay and indicates the bandwidth at which magnetomicrometry can track the muscle tissue length. The data are shown as a stacked histogram, with blue, orange, and purple data corresponding to Birds A, B, and C, respectively. Data are from all turkey gait cycles used to compare magnetomicrometry against fluoromicrometry. The 99th percentile time delay (Δ_t99%_) is labeled for each bird. The ninety-ninth percentile time delay for all birds was less than one millisecond.

**Supplementary Figure 6:**
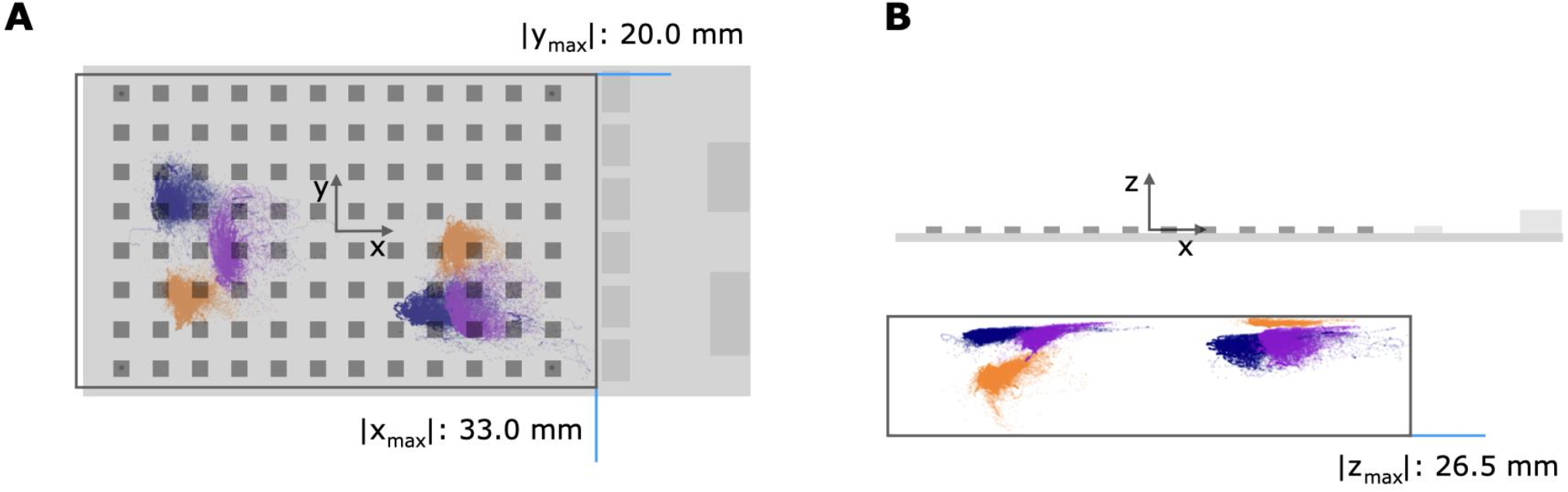
Extent of Bead Positions During Ramp Ascent and Descent and Vertical Ascent and Descent. Analysis of the magnetic bead position data (shown) recorded during all variable terrain activities revealed the extent of the magnetic bead positions during the tracking. We used these bead position maximums to design a benchtop test (see supplementary Figure 7) to verify the validity of our magnetomicrometry measurements during the variable terrain activities.

**Supplementary Figure 7:**
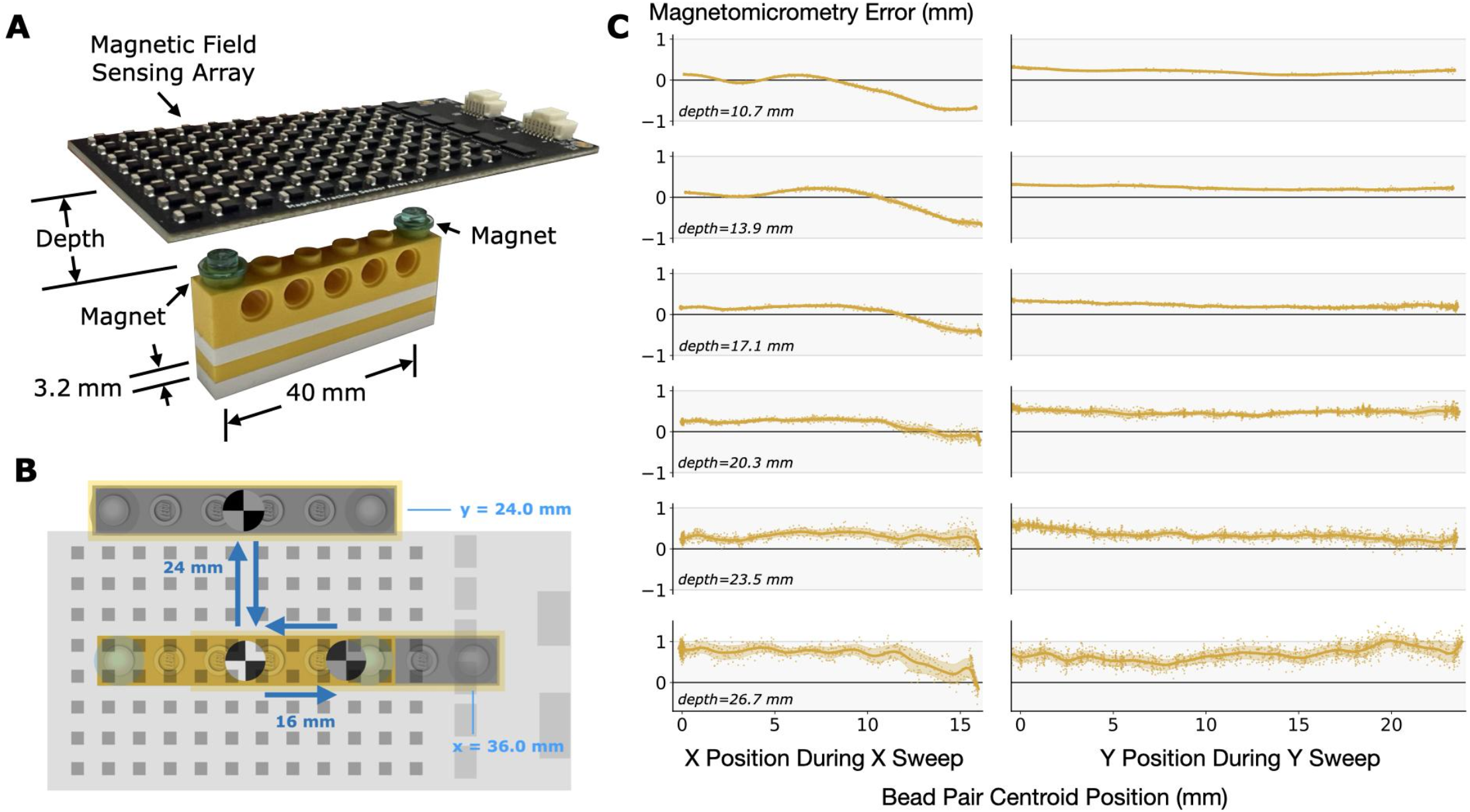
Benchtop Magnetomicrometry Validation. **(A)** Two magnets were placed 40 mm apart in a 1×6 LEGO Technic block and centered under the sensing array. We used the tracked magnet z-position as a guide in setting up the minimum depth measurements, and we enforced the remaining depths by adding/removing 3.2-mm-thick 1×6 plates under the Technic block. **(B)** We manually swept the magnets out and back to center along the x and z axes. We set the sweep trajectory to sweep just beyond the volume within which the magnets were tracked during the variable terrain activities (see Supplementary Figure 6). For reference, the centroid of the two beads is labeled at the origin and at the extent of the sweep trajectories. **(C)** The vertical axis represents the magnetomicrometry error, and the horizontal axes represent the centroid x and y position for the two sweep trajectories at each depth. The submillimeter error range is marked with a gray background. A maximum error of 1.463 mm was found at a test location just beyond the tracked bead position extent (bottom right plot).

